# Listening to classical music influences brain connectivity in post-stroke aphasia: a pilot study

**DOI:** 10.1101/2023.09.27.559699

**Authors:** Maryane Chea, Amina Ben Salah, Monica N. Toba, Ryan Zeineldin, Brigitte Kaufmann, Agnès Weill-Chounlamountry, Lionel Naccache, Eléonore Bayen, Paolo Bartolomeo

## Abstract

Stroke-induced aphasia is a leading cause of cognitive disability. The healthy right hemisphere may play a role in aphasia compensation. Music-based therapy, known to enhance cognitive functions after stroke, offers a potential intervention due to its impact on brain connectivity, engaging both hemispheres.

In this proof-of-concept study, we aimed to assess the feasibility and effectiveness of music-assisted therapy for language disorders following left hemisphere strokes. We also sought preliminary insights into the impact of music on post-stroke brain connectivity. We enrolled four right-handed patients (one female and three males, mean age 57.75 years, SD 7.63) who experienced their first left-hemisphere stroke inducing chronic aphasia symptoms three months post-stroke. Patients were randomly assigned to receive either two weeks of music therapy, involving daily 2-hour sessions of listening to instrumental music by Haydn, Mozart, and Beethoven, in addition to standard rehabilitation, or two weeks of standard care, using a crossover design. Patients underwent cognitive and neuroimaging assessments at baseline, crossover, and study end, with preliminary evaluation one week before the study to confirm stable deficits.

All patients listened to the same curated playlist, accompanied by narrations situating the music in its with historical context. Cognitive assessments included the Core Assessment of Language Processing (CALAP) and a French aphasia battery (Bilan Informatise d’Aphasie, BIA). EEG and MRI data were analyzed, focusing on functional connectivity metrics (weighted symbolic mutual information, permutation entropy) and white matter tractography. Preliminary findings indicate that music therapy may be a feasible and promising approach to aphasia rehabilitation, with improvements observed in language tests and brain connectivity metrics.

Despite limitations such as a small sample size and patient heterogeneity, these preliminary results suggest that intensive classical music listening may enhance language abilities in post-stroke aphasia patients. Additionally, using EEG connectivity metrics originally designed for non-communicative patients provides a novel avenue for monitoring brain plasticity during rehabilitation.

---

Dear Editor,

Stroke-induced aphasia is a leading cause of acquired disability worldwide. Some patients may partially recover, but they do not often retrieve their full previous cognitive abilities. Traditionally, post-stroke cognitive deficits were considered as resulting from focal cortical dysfunction. Nevertheless, abundant evidence now emphasizes the importance of disruption of activity in highly distributed, large-scale neural networks, connected by long-range white matter tracts [1]. Evidence is, however, conflicting on the role of activity in the right, non-damaged hemisphere in recovery from aphasia. Some studies suggest that it has a negative impact, while others suggest that it serves an adaptive, compensatory function [2,3]. Viable interhemispheric connectivity may be important in promoting a shift in activity in the healthy hemisphere from maladaptive to compensatory [4]. If this is the case, then encouraging inter-hemispheric integration may be a helpful approach to facilitate recovery from post-stroke aphasia. A possible candidate intervention could be listening to music [5], which has been shown to improve cognitive functions after stroke [6]. The beneficial effect of music might be due to improved brain connectivity, as music processing typically involves coordinated bi-hemispheric activity in neurotypical individuals [7,8].

The goals of this proof-of-concept study were to explore the feasibility and effectiveness of using music-assisted therapy to treat language disorders following left hemisphere strokes, and to obtain preliminary data on the effect of music on brain connectivity after stroke. Inclusion criteria were a first left unilateral stroke inducing chronic signs of aphasia 3 months post-stroke. Exclusion criteria were impaired vigilance, confusion, general mental deterioration, psychiatric disorders, prior history of neurological disease, or contraindications to MRI. Four right-handed patients with a first left-hemisphere stroke (one woman and three men; mean age 57.75 ± 7.63 years) were included from May to July 2022 (see Additional Table 1 for patients’ demographic and clinical characteristics). Patients 1-2 had signs of non-fluent aphasia and Patients 3-4 had very mild signs of fluent aphasia. All patients gave their written informed consent according to the Declaration of Helsinki. The study was promoted by INSERM and approved by the IRB Ile-de-France I. Each patient was assessed multiple times, thus serving as his/her own control. This allowed us to conduct individual analyses using a multiple single-case experimental design [9]. Patients were randomly assigned to receive either 2 weeks of therapy involving listening to instrumental music by Haydn, Mozart, and Beethoven for 2 hours per day in addition to their usual rehabilitation, or 2 weeks of standard care, in a crossover design (Figure 1). This means that the patients who received the music therapy in the first two weeks of the study only received standard care in the second two weeks, and vice versa, allowing us to compare the effects of the two treatment conditions. Music was presented in high quality audio-video format. All patients listened to the same playlist, which was created in partnership with professional musicians. To optimize the participants’ engagement, short narrations were provided before each musical excerpt to provide historical context and explain the composer’s intended message. There was a 15-minute break after the first hour of music. Classical music from the Vienna school was selected because it strikes a good balance between expected and unexpected elements, and the length of the typical musical phrases is well-suited to engage working memory, which could help cognitive recovery [5].

**Figure 1:**
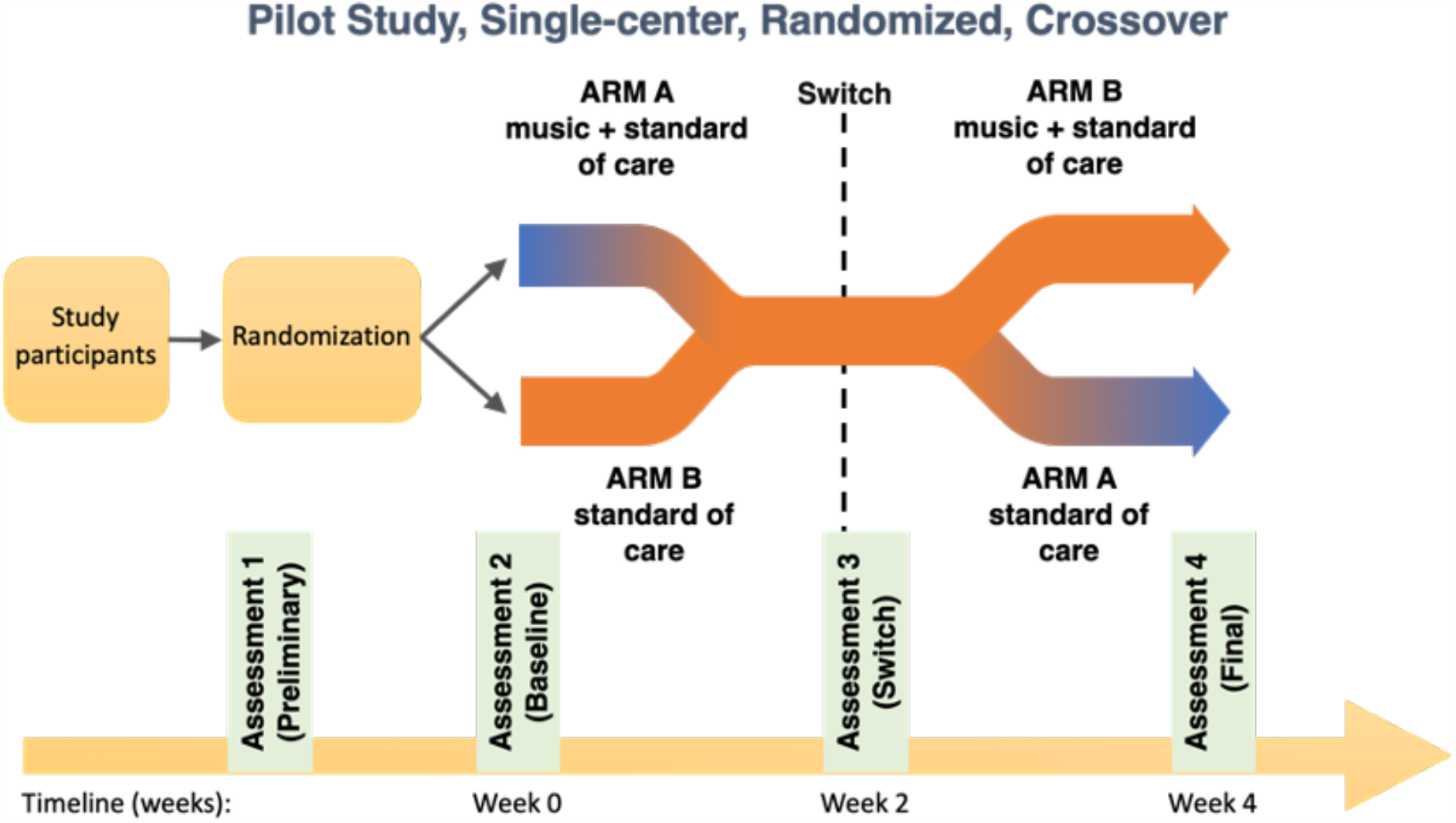
Timeline of the protocol. During Assessment 1, clinical evaluation was performed twice so as to verify stability of language impairment. Assessments 2 to 4 were both clinical (language tests) and with imaging (EEG, MRI). Switch between arms was performed after 14 days.

Cognitive assessment and neuroimaging were conducted at baseline, at crossover and at the end of the study. In addition, a preliminary cognitive assessment was conducted one week before the start of the study to confirm that the deficit was stable over time. Language was assessed using the Core Assessment of Language Processing (CALAP) screening test [10], and a French aphasia battery (Bilan Informatisé d’Aphasie, BIA) [11], which offers a more detailed assessment. We conducted McNemar tests for total and item scores on CALAP and BIA, with continuity correction where appropriate. The alpha level was set at 0.05.

EEG and MRI were obtained using methods previously described by our team [12, 13]. Task-free, high-density EEG recordings were sampled at 250 Hz with a 256-channel geodesic sponge sensor net referenced to the vertex. To assess functional connectivity, we determined the weighted symbolic mutual information metric (wSMI) of resting state EEG theta frequency band [14]. This metric captures the nonlinear coupling between pairs of electrodes and quantifies the amount of information shared between the electrodes. We also calculated permutation entropy (PE) in the theta band [14] to assess complexity.

MRI scans, which we were able to obtain only for Patient 3, were acquired with a 3.0 Tesla Siemens MRI and included a diffusion tensor (DT) sequence. During the preprocessing, eddy current-induced distortions were removed and motion distortion corrections were computed before the estimation of the diffusion tensors. Using standard computational algorithms, fractional anisotropy (FA) was calculated in the native space and white matter (WM) tractography was then performed by following a region-of-interest (ROI) approach [13]. Specifically, on the basis of previous work on tractography in stroke patients [15–17], we considered the following white matter bundles: arcuate fasciculus (AF), inferior longitudinal fasciculus (ILF), inferior fronto-occipital fasciculus (IFOF), fronto-parietal superior longitudinal fasciculus (SLF), frontal aslant tract, and corpus callosum (see Additional material).

Neuropsychological results showed no significant change in cognitive functions between the preliminary and baseline assessments in any of the patients (see Additional material), indicating that their aphasia was stable at inclusion (Figure 2). Patients 3-4 obtained near-normal scores on the aphasia batteries at inclusion; they were included as control patients to assess the feasibility of the procedure. After two weeks of listening to music, Patient 1 (Arm A) and Patient 2 (Arm B) obtained increased BIA scores by 21.31% (corrected McNemar’s □^2^, 9.00, p=0.003 and 9.10% (uncorrected □^2^, 7.53, p=0.006), respectively. In contrast, after receiving standard rehabilitation, Patients 1 and 2 did not show any significant change in test performance. In Patients 3 and 4, language performance was near ceiling at inclusion, and did not change significantly after either music-assisted or standard rehabilitation.

**Figure 2:**
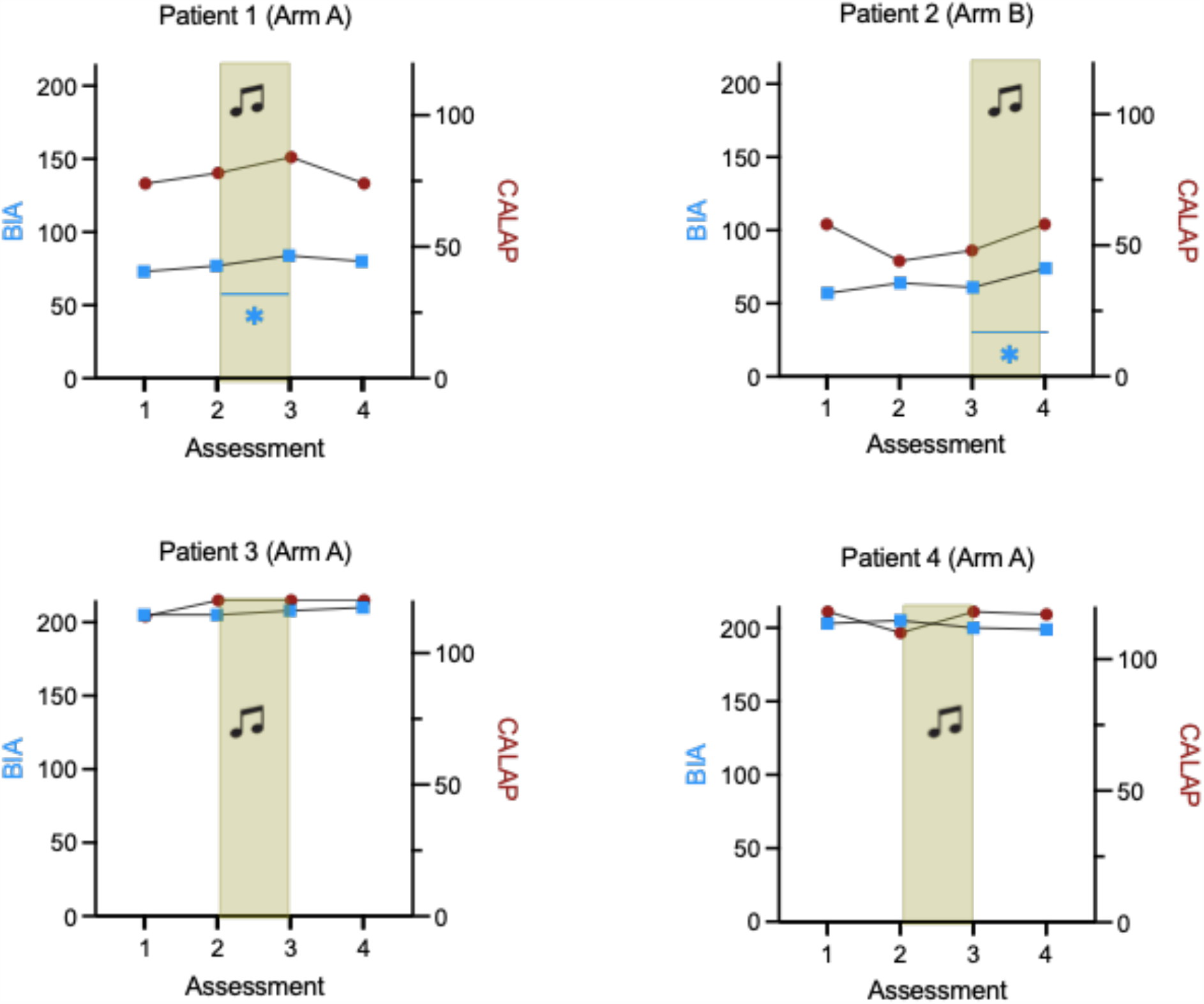
Patients’ performance on language tests over the 4 assessments. Core Assessment of language processing (CALAP) (max 120) and Batterie Informatisée Aphasie (BIA) scores (max 215) in arm A and arm B patients before and after the music protocol (in color, the 2 weeks of listening to music). * McNemar test, p < 0.05

**Figure 3:**
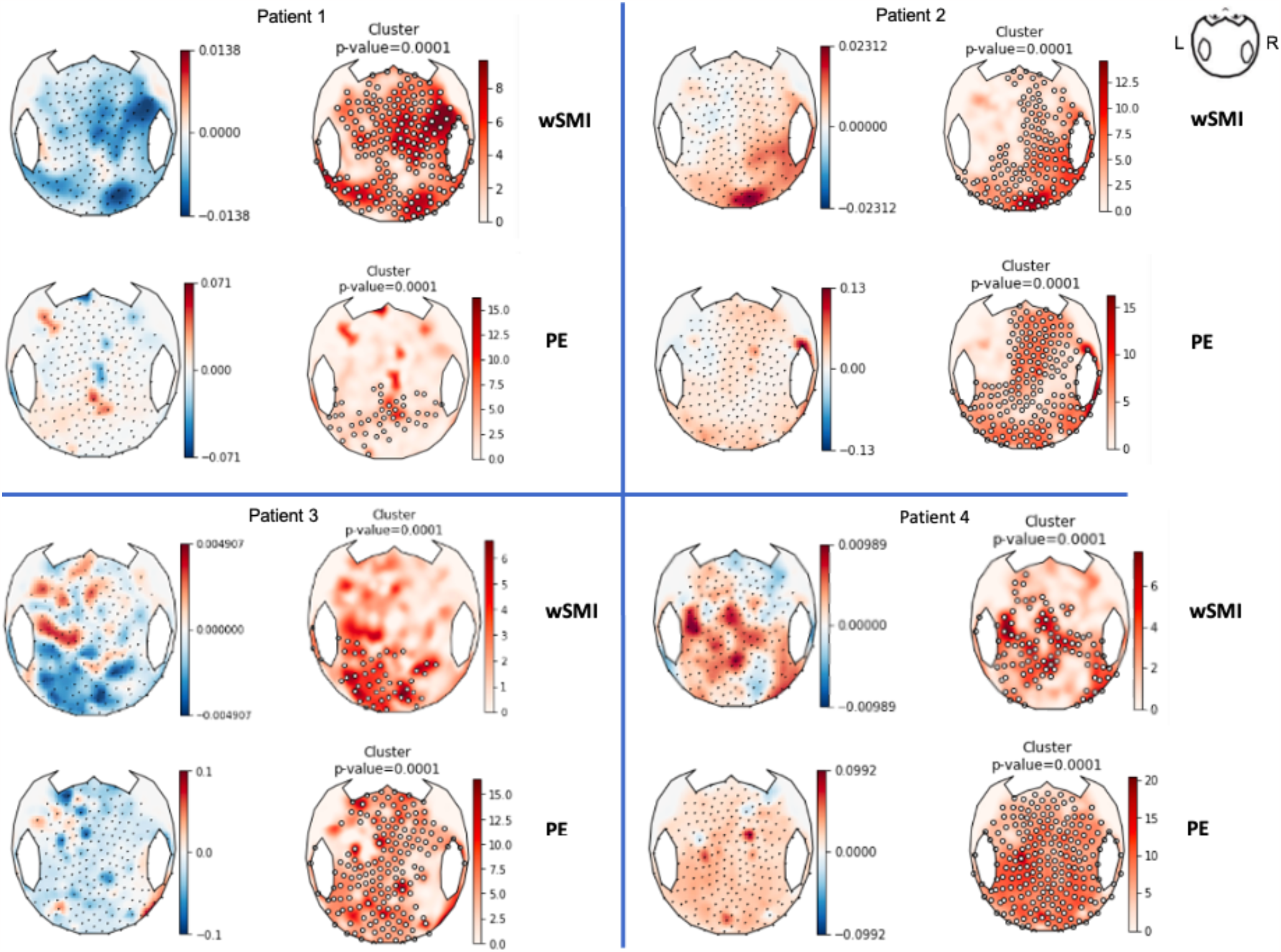
EEG Functional connectivity before and after 2-week music listening. For each patient (1 to 4), comparison of weighted symbolic mutual information (wSMI) and permutation entropy (PE) before and after 2 weeks of music listening. A t-test was performed to compare values before and after music listening. Clusters are represented (p = 0.0001).

Regarding EEG connectivity, Patient 1’s EEG was difficult to interpret due to artifacts on the temporal regions, perhaps as a consequence of hemicraniectomy. After music-assisted rehabilitation, Patient 2 showed an increase in the theta wSMI in the parieto-occipital region of the right, non-damaged hemisphere, along with an increase in permutation entropy (PE) (t-test: p = 0.0001, cluster p = 0.0001 for both), reflecting an increase in signal complexity/informativity. For Patients 3 and 4, an increase in wSMI and PE was observed in the left, lesioned hemisphere after music-assisted rehabilitation. After standard rehabilitation, an increase in theta wSMI and PE was found in the centro-frontal and parietal regions of the left hemisphere (t-test: p = 0.00001, cluster p = 0.0001).

Patients 1, 2 and 4 declined to complete the lengthy MRI sessions. In the tractography analysis, after two weeks of music therapy, Patient 3 (the only patient for whom a complete MRI follow-up was available) showed an increase in FA in bilateral IFOFs, corpus callosum (particularly the tapetum), the right hemisphere SLF III, and the right frontal aslant tract (Additional Table 3). At the final assessment, FA in these bundles had decreased compared to the post-music assessment, although it remained higher than the baseline value before music therapy, potentially indicating neuroplasticity. There was no significant change in FA values in the SLF I and II networks.

These pilot results suggest that intensive listening to classical music is a feasible and potentially beneficial approach for aphasia rehabilitation. Patients 1 and 2 showed some improvement in language tests after listening to music. In Patient 2, music, but not standard rehabilitation, was associated with cognitive improvement and increased EEG connectivity in the non-lesioned hemisphere. Patients 3 and 4 had very mild language deficits and therefore less margin for cognitive improvement. However, in both these patients listening to music was followed by increases in EEG metrics of brain connectivity in the left hemisphere. Moreover, in Patient 3 music-assisted rehabilitation was followed by changes in white matter microstructural parameters, suggesting an increase in interhemispheric (corpus callosum) and intrahemispheric (IFOF, SLF III, frontal aslant) connectivity. This result is in line with previous research [18], suggesting that instrumental music may lead to broader changes in connectivity in the post-stroke brain, as opposed to listening to vocal music which primarily stimulates the left hemisphere.

In terms of feasibility, our music-assisted intervention was well-received by all patients, even though none of them had prior musical training. Using a limited selection of classical music did not hinder the patients’ adherence to the therapy protocol, which proved to be a relatively inexpensive and non-tiring form of treatment. The cumulative duration of music sessions in this study was 20 hours, which falls within the range of previous music-assisted rehabilitation studies (6-30 hours [19]).

The primary limitation of this proof-of-concept study is the small sample size and the heterogeneity of patients. Also, clinical and neuroimaging evaluations were not performed blind to the patient’s arm. There was a difference between the control and experimental condition in the social aspects, as patients listening to music did so in small groups rather than individually. This choice was made to maximize the potential benefits of music. Although it introduces the possibility of a confounding effect, we believe that the potential clinical advantages outweigh this risk. Any potential confound can be addressed in future studies.

Despite these limitations, our initial findings are promising regarding the feasibility of the procedure and the potential for enhancing language abilities in post-stroke aphasia, through listening to classical instrumental music. The application of wSMI theta and PE, which are EEG connectivity metrics originally designed for noncommunicative patients [12], to analyze and follow-up brain connectivity after focal strokes, represents a novel and promising approach to monitor brain plasticity changes during rehabilitation.

## Data Availability Statement

The datasets used and/or analyzed during the current study are available from the first author on reasonable request.

## Acknowledgments

The authors would like to thank the patients and their relatives for their patience and cooperation, Maestro Lorenzo Coppola and the participating orchestras (notably the Paris Mozart Orchestra), the CENIR neuroimaging team at Paris Brain Institute (notably Benoît Béranger), Alexandre Guerre, Laura Vincent, Jean-Jacques Repain and Yannick Mahé at Sorbonne University.

