## Additional material for "Listening to classical music influences brain connectivity in post-stroke aphasia: a pilot study"

**Single-item assessment for BIA and CALAP**

Here and elsewhere [1, 2], we assessed patients’ performance changes by using McNemar 𝜒², with continuity correction when before-after pairs were less than 10. After 2 weeks of music listening, the item-by-item analysis of Patient 1’s performance showed that only BIA pseudoword repetition showed a significant improvement (corrected McNemar 𝜒² 4.16 with correction, p = 0.04). For Patient 2, no BIA item-by-item assessment emerged as significant.

After standard rehabilitation, there were no significant changes for any single items.

**Additional Table 1. Patients’ clinical and demographic characteristics**

| **Patient** | **Age** | **Sex** | **Laterality** | **Days after stroke** | **Etiology** | **Aphasia** | **Education (years)** | **NIHSS**  **at inclusion** | **Motor/ sensory disorders** |
| --- | --- | --- | --- | --- | --- | --- | --- | --- | --- |
| 1 | 60 | M | R | 595 | I | TMA | 20 | 14 | Y/Y |
| 2 | 65 | M | R | 129 | H | TMA | 23 | 13 | Y/Y |
| 3 | 59 | M | R | 89 | I | fluent | 18 | 1 | Y/Y |
| 4 | 47 | F | R | 88 | I | fluent | 18 | 1 | N/N |

F : frontal lobe, T: temporal lobe, O: occipital lobe, P : parietal lobe, I : insula, TMA : transcortical motor aphasia, R : right, I : ischemic, H : hemorrhagic, Y: yes, N: no; M : male, F: female.

**Additional Figure 1. Lesional profiles of patients included in the study. Arrows indicate the presence of lesions. L: left**

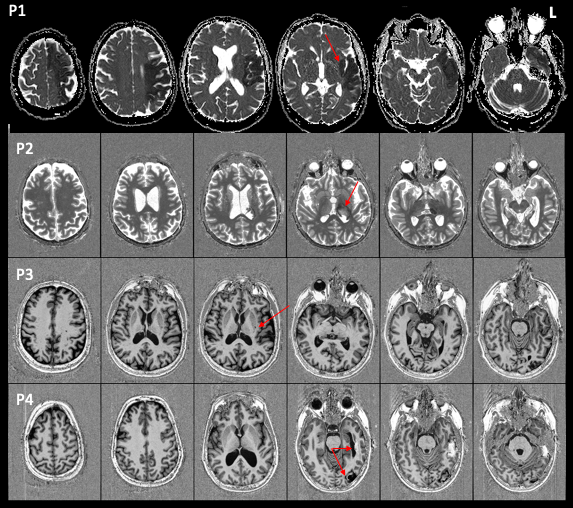

**Additional Table 2. Individual test changes**

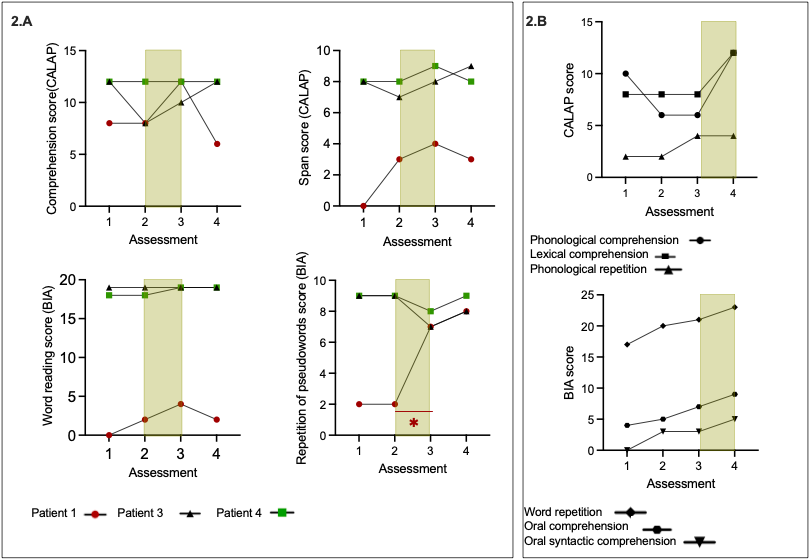

Fig 2. A: Results of CALAP and BIA for Arm A (2.A, n = 3) and Arm B (2.B, n = 1), before and after two weeks of music protocol. The maximum value on each y-axis corresponds to the highest possible score that can be awarded for the respective item.

**Additional Table 3. Normalized metrics of MRI structural connectivity**

| **Assessment** | **IFOF L** | **IFOF R** | **Corpus Callosum Forceps Minor** | **Corpus Callosum**  **Corps** | **Corpus Callosum**  **tapetum** | **Corpus callosum**  **forceps major** | **Arcuate**  **fasciculus L** |
| --- | --- | --- | --- | --- | --- | --- | --- |
| FAn-V1 | 1.12 | 1.09 | 1.03 | 1.21 | 1.22 | 1.24 | 1.09 |
| **FAn-V2** | **1.15** | **1.12** | **1.09** | 1.22 | **1.38** | **1.31** | 1.08 |
| FAn-V3 | 1.13 | 1.10 | 1.04 | 1.22 | 1.31 | 1.34 | 1.09 |
| **Assessment** | **Arcuate fasciculus R** | **SLF I L** | **SLF I R** | **SLF II L** | **SLF II R** | **SLF III L** | **SLF III R** |
| FAn-V1 | 1.09 | 0.38 | 0.41 | 0.92 | 0.93 | 0.97 | 0.93 |
| **FAn-V2** | 1.07 | 0.38 | 0.41 | 0.91 | 0.93 | 0.97 | **0.96** |
| FAn-V3 | 1.05 | 0.38 | 0.41 | 0.93 | 0.92 | 0.97 | 0.94 |

Evolution of normalized fractional anisotropy (FAn) characterizing MRI-derived white matter fibers diffusion tensor of Patient 3, according to the sessions (V1: before listening, V2: at the end of the 2 weeks music + traditional rehabilitation, V3: 4 weeks after the beginning of the protocol). FAn: normalized fractional anisotropy, L: left, R: right, IFOF: inferior fronto-occipital fasciculus, SLF: superior longitudinal fasciculus. In bold are indicated the FAn values that increased between the baseline and post-music evaluation.

| **Assessment** | **IFOF L** | **IFOF R** | **Corpus callosum Forceps Minor** | **Corpus callosum**  **Corps** | **Corpus call**  **tapetum** | **Corpus callosum forceps major** | **Arcuate**  **fasciculus L** | **Frontal Aslant**  **L** |
| --- | --- | --- | --- | --- | --- | --- | --- | --- |
| nRD-V1 | 0.97 | 0.98 | 1.06 | 0.89 | 1.37 | 1.00 | 0.89 | 1.01 |
| **nRD-V2** | **0.94** | 0.98 | **1.02** | **0.88** | **1.12** | **0.94** | 0.90 | 1.01 |
| nRD-V3 | 0.95 | 0.97 | 1.06 | 0.89 | 1.20 | 0.96 | 0.90 | 1.01 |
| **Assessment** | **Arcuate fasciculus R** | **SLF I L** | **SLF I R** | **SLF II L** | **SLF II R** | **SLF III L** | **SLF III R** | **Frontal Aslant R** |
| nRD-V1 | 0.91 | 1.02 | 0.99 | 1.01 | 1.02 | 0.99 | 1.01 | 1.02 |
| **nRD-V2** | 0.92 | 1.02 | 0.97 | 1.03 | 1.02 | 0.99 | **0.99** | **1.00** |
| nRD-V3 | 0.94 | 1.02 | 0.98 | 1.01 | 1.03 | 0.99 | 1.01 | 1.02 |

Evolution of normalized radial diffusivity (nRD) in diffusion sequence, characterizing MRI-derived white matter fibers diffusion tensor of Patient 3, according to the sessions (V1: before listening, V2: at the end of the 2 weeks music + traditional rehabilitation, V3: 4 weeks after the beginning of the protocol). L: left, R: right, IFOF: inferior fronto-occipital fasciculus, SLF: superior longitudinal fasciculus. In bold are indicated the FAn values that increased between the baseline and post-music evaluation.

**Additional figure 4. EEG functional connectivity before and after traditional rehabilitation**

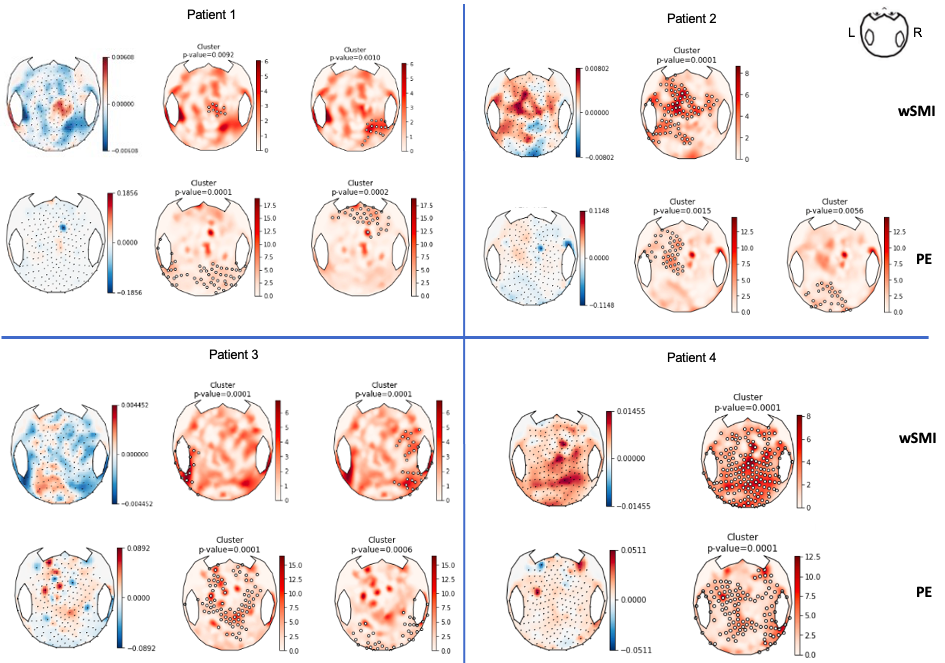

Weighted symbolic mutual information (wSMI) and permutation entropy (PE) before and after 2 weeks of standard rehabilitation for each patient (1-4). T-tests were performed to compare values before and after music listening. Cluster permutation analysis on EEG data, based on t-test, are represented (p < 0.05). Scalp topographies show the statistic T-stat, comparing before and after 2 weeks of music. Thick electrodes show presence of an observed effect in cluster-permutation analysis.

**Methods**

*Clinical Assessment*

In order to assess the stability of the cognitive sequelae of the stroke, each subject was assessed twice before inclusion, thus providing a baseline assessment. NIHSS was assessed at baseline.

In this protocol, we evaluated oral language through oral expression (oral naming, repetition), oral comprehension (naming pictures, semantic matching, syntactic comprehension), written comprehension (naming written words), and working memory (span).

*EEG preprocessing and analysis*

All analyses were then performed at the group level, using the mean of the individuals in the group and comparing the pre-intervention means to the post-intervention means, using a paired univariate analysis and then using a two-step permutation approach based on spatial clusters. All markers were first computed at the subject level: a value was obtained for each epoch at each channel (or channel pairs for wSMI). The values were then averaged over the epochs to obtain a two-dimensional topographic representation over the 224 electrodes for each subject.

*EEG acquisition*

Resting EEG recordings, lasting 10 minutes, with eyes closed, in a dark, quiet room, were recorded at a sampling rate of 250 Hz using a NetAmps 300 amplifier (Electrical Geodesics) with a 256-channel geodesic sensor net referenced to the top. Impedances were set below 100 kΩ.

*EEG pre-processing*

EEG recordings were bandpass filtered (using a 6th-order Butterworth high-pass filter at 0.5 Hz and an 8th-order Butterworth low-pass filter at 45 Hz) with 50 Hz and 100 Hz notch filters. Channels that exceeded an amplitude of 150 μV in more than 50% of the epochs were rejected. The remaining epochs were digitally transformed to an average reference. The rejected channels were interpolated. EEGs were considered to pass this preprocessing step if at least 75% of channels and at least 30% of epochs were retained.

*Quantitative markers studied*

The quantitative markers chosen are derived from resting EEG recordings following previous methodological methods [14, 15] and have been used to measure the effect of an intervention on the level of consciousness in non-communicating patients by comparing these measurements before and after intervention [16].

- Spectral domain: the density of the power spectrum in each frequency band (δ: 1-4 Hz; θ: 4-8 Hz; α: 8-12 Hz; β: 12-30 Hz; γ: 30-45 Hz) was calculated by fast Fourier transformation with Welch's method with a periodogram of 512 ms and an overlap of 400 ms. Raw and normalized spectral power (the sum of the power in one frequency band related to the power over all frequency bands in the spectrum sum) are reported for each frequency band.

- Connectivity: Functional connectivity was assessed using weighted symbolic mutual information (wSMI). This metric, capable of capturing the nonlinear coupling between pairs of electrodes, was introduced by King et al [17]. It quantifies the amount of information shared between the electrodes in each frequency band.

- Complexity: We measured the permutation entropy in the theta band, which has been successfully applied to EEG detection of loss of consciousness under anesthesia [18], but also more recently in studies of neurodegenerative diseases [19].

All markers were first computed at the subject level: a value was obtained for each epoch at each channel (or pairs of channels for the wSMI). Values were then averaged across epochs to obtain a 2-dimensional topographic representation across all 224 electrodes for each subject.

We also measured the PE in the theta band to study complexity. All markers were first calculated at the subject level: a value was obtained for each epoch at each channel (or channel pairs for wSMI). Values were then averaged across epochs using the 80% adjusted mean (average of the distribution after eliminating the lowest 10% and highest 10% values) to obtain a two-dimensional topographic representation across all 224 electrodes for each subject. All analyses were then performed at the group level, using the mean of the individuals in the group and comparing the pre-intervention means to the post-intervention means, using a paired univariate analysis and then using a two-step permutation approach based on spatial clusters.

*Tractography study* DTI images were acquired on a 3.0 Tesla Siemens MRI at the CENIR, with a 64-channel head coil. For diffusion imaging, we used echo planar sequences (EPI) with 98 directions (b = 3000 sec/mm2, TR/TE = 3370 ms TE 89.20 ms, flip angle = 78 degrees, FOV = 210 mm, matrix size = 140 X 140 X 96, voxel size = 1.5 mm, slice thickness = 1.50 mm). Processing and visualization (FA, RD, tract-graphs) were done with DSI-Studio (<https://dsi-studio.labsolver.org/>). Preprocessing was performed using FSL (v. 6.0.3) and Explore DTI. It included steps to correct artifacts, head movements andeddy currents.. We performed quality control per slice with DSI Studio.

The tractography analyses were performed with DSI Studio. The accuracy of b-table orientation was examined by comparing fiber orientations with those of a population-averaged template [7]. The restricted diffusion was quantified using restricted diffusion imaging [8]. The diffusion data were reconstructed using generalized q-sampling imaging [9] with a diffusion sampling length ratio of 1.25. A deterministic fiber tracking algorithm [10] was used with augmented tracking strategies [11] to improve reproducibility. The anatomy prior of a tractography atlas [12] was used to map tracts with a distance tolerance of 16 mm in the ICBM152 space. The anisotropy threshold, the angular threshold (from 15 degrees to 90 degrees) and the step size (from 0.5 voxel to 1.5 voxels) were randomly selected. Tracks with length shorter than 20 or longer than 300 mm were discarded. Topology-informed pruning [13] was applied to the tractography with 16 iterations to remove false connections. Fractional anisotropy (FA) and radial diffusivity (RD) of selected bundles were analyzed between different evaluation times.

We focused on the white matter bundles that are involved in language. In the dual pathway model [3], the main centers are connected by white matter tracts. We distinguish the bilaterally distributed ventral network involved in language comprehension and the dorsal network, predominantly on the left, important for production. In the dorsal network, the inferior frontal gyrus, the anterior and posterior portions of the insula, the precentral gyrus, and the supramarginal gyrus are connected by the arcuate fasciculus (AF) and the posterior components of the superior longitudinal fasciculus (SLF). The ventral network, including the inferior fronto-occipital fasciculus (IFOF), the inferior longitudinal fasciculus (ILF), the external capsule, and the uncinate fasciculus, connects the superior temporal gyrus, the superior temporal sulcus, the middle and inferior temporal gyri, and the anterior temporal lobe.
